# Modelling the effects of bioclimatic characteristics and climate change on the potential distribution of a monospecific species *Colophospermum mopane* (Benth.) Léonard in southern Africa

**DOI:** 10.1101/2021.06.03.446954

**Authors:** Boniface K. Ngarega, Valerie Farai Masocha, Harald Schneider

**Affiliations:** Wuhan Botanical Garden, Chinese Academy of Sciences, Wuhan, China; Centre for Integrative Conservation, Xishuangbanna Tropical Botanical Garden, Chinese Academy of Sciences, Menglun, China

**Keywords:** Climate change, Southern Africa, Species distribution modelling, *Colophospermum mopane*, Maxent

## Abstract

Global climate change is gradually changing species distribution and their patterns of diversity. Yet, factors that influence the local distribution and habitat preferences for southern African species remain largely unexplored. Here, we computed the suitable habitats in the southern African region for *Colophospermum mopane* (Benth.) using the maximum entropy (Maxent) modeling approach. We utilized one Global Circulation Model (GCM) and three Representative concentration pathways (RCPs) to determine the current and future distribution of *C. mopane*. The results showed that the distribution of *C. mopane* was mainly influenced by solar radiation, annual temperature range, and annual precipitation. According to the species response curves, this species preferred habitats with annual precipitation of 130-200 mm, an annual temperature range of 28° C, and elevations of about 500 m above sea level. The results highlight that the geographic range of *C. mopane* is likely to expand along the borders of Zambia and Zimbabwe in the future, particularly in the miombo plains. Conversely, suitable habitat areas reduce significantly in the eastern area of the southern African region, while the western areas expand. Overall, the appropriate habitat areas will likely decline in the 2050s under both RCPs and expand in the 2070s under the two scenarios. This knowledge is important for landscape planners and rangeland managers working to safeguard biodiversity from extinction.

**Highlights:**

- High reliability of models in habitat suitability modelling for *C. mopane*
- Solar radiation is the most significant variable for the current distribution of mopane.
- Climate change is and will reduce the habitat suitability of our target species.

## 1. Introduction

*Colophospermum mopane* (Benth.) Léonard, typically known as the Southern African Mopane, is a monospecific woodland species belonging to the family Fabaceae (Moura, 2017). This plant species is extensively distributed in the hot, low-lying areas of southern Africa’s savannas, covering regions of about seven countries (Fig 1) (Burgess et al., 2004; Maquia et al., 2019). This dominant tree or multi-stemmed shrub in the mopane woodlands is globally considered ecosystems with irreplaceable species endemism, making it biologically unique. Being a predominant native species, it forms the most important socio-economic and environmental vegetation in the area and a ready source of indigenous woody products, food, and medicine for a large number of urban and rural residents highly reliant on these ecosystems (Mojeremane & Lumbile, 2005; Dewees et al., 2010; Rosenstock et al., 2019). As it is suited to a wide range of soil and microclimates, the plant takes on a variety of growth types, ranging from shrub-like to tall, slender trees with stunning leaf canopy. Its leaves are a primary habitat for the caterpillar worms (*Gonimbrasia belina*), which forms a significant feed, and income source to the region, particularly the rural communities (Mojeremane & Lumbile, 2005; Langley et al., 2020).

**Figure 1.**
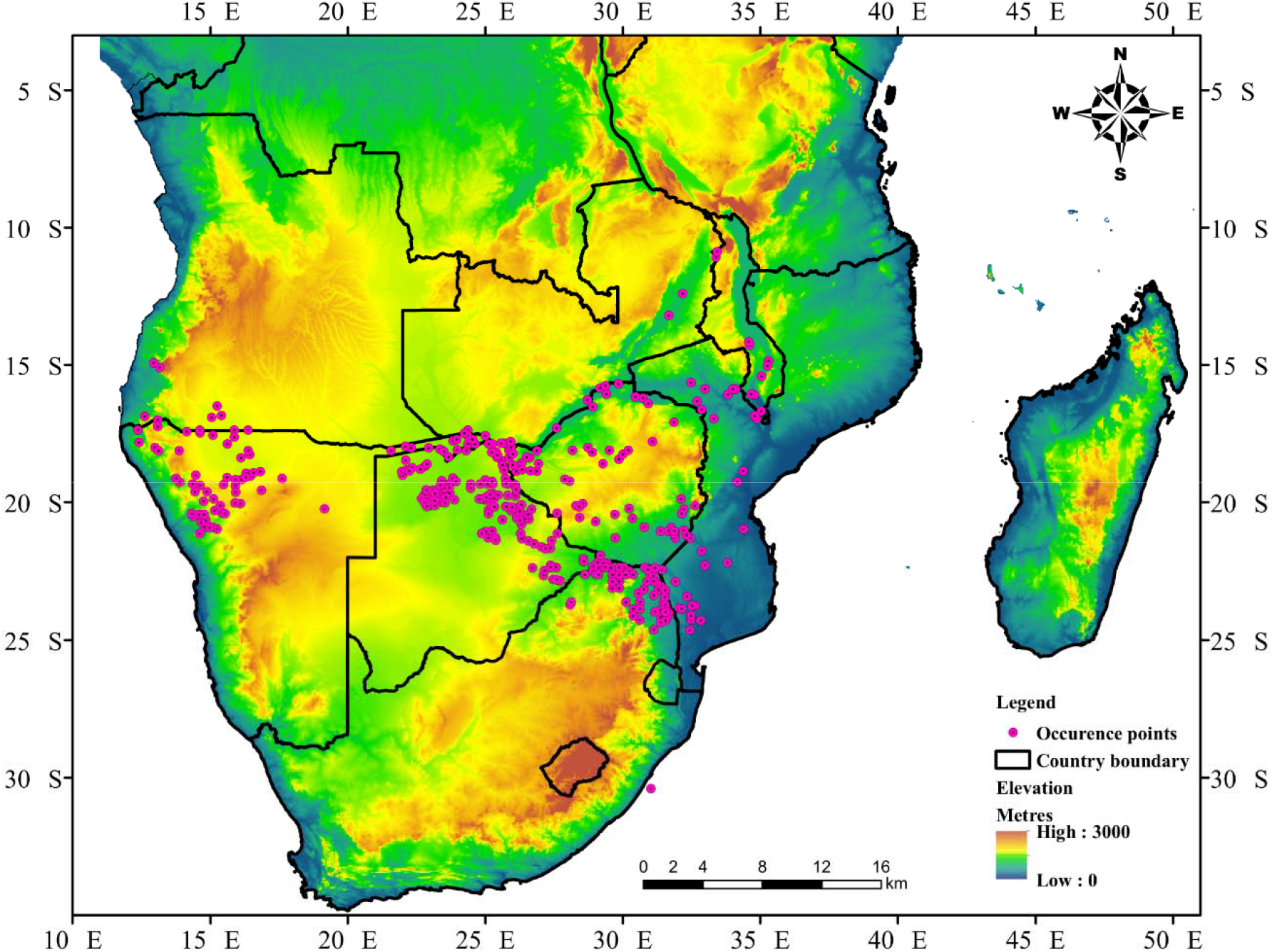
Localities and occurrence records of *C. mopane* obtained from the GBIF and Zambia Flora database.

Mopane is thought to display gregariousness by suppressing other woody plants through various mechanisms, including the release of allelopathic compounds (Daru et al., 2016), making it a promising candidate for investigating landscape genomics. In addition, Mopane forms part of biodiversity that has a global impact on water carbon sequestration, as well as energy and water balances (Handa et al., 2020; Mlambo et al., 2005). Furthermore, the ecological dynamics of *C. mopane* have been reported to be considerably influenced by a combination of climate change and non-climatic factors such as fire (Kennedy and Potgieter, 2003), day length, and animal disturbances (Stevens et al., 2018, 2014). However, its ecological niche is experiencing human population expansion, which consequently exerts pressure on the woodlands through activities such as mining, farming, and clearing for infrastructure. With the change of climate, potentially increased and recurring drought seasons are expected in most parts of the world, resulting in desertification (Krug, 2017). Therefore, the distribution modelling and future prediction of drought-tolerant vegetation species such as Mopane, becomes a necessity.

Climate is often related to the plants’ global distribution because it reflects the availability of moisture and energy for plant growth, and therefore, forecasting species distribution is an important tool, particularly for ecologists and conservationists, in mitigating climate change. Statistical species distribution models (SDMs) have been broadly used to forecast species distributions’ potential changes under climate change (Austin and Van Niel, 2011; Booth, 2018). SDMs relate environmental variables to well-known species occurrence locations to establish abiotic conditions under which organisms will survive (Guisan & Thuiller 2005). Bioclimatic constructions are well known as the best, if not the only, method to model species distribution in response to climate (Longmore and Busby, 1986). However, some recent research has indicated that non-climatic environmental factors such as fire (Kennedy and Potgieter, 2003; Lewis et al., 2017; Makhado et al., 2014), and day length (Dallas et al., 2017), can affect the distribution of plants.

Climate change has had a major effect on natural and human environments in recent decades. These effects of climate change, regardless of their origin, demonstrate the resilience of natural and human environments to changes in climatic systems’ structure, interference between their elements, or changes in external factors, either spontaneously or due to anthropogenic causes (Ipcc, 2014). Notably, researchers and wildlife managers place a high emphasis on factors that impact species distributions and habitat selection. As such, it is vital to understand the impact of variables on species occurrence (Baldwin, 2009).

Many species’ future distributions are increasingly being mapped using (SDMs) and ecological niche models (ENMs) (Fourcade et al., 2014). The purpose of a modeling approach is to estimate the ecological appropriateness of the ecosystem according to environmental variables (Phillips, 2008., Blanco et al., 2020). In this study, we opted to predict the distribution patterns of *C. mopane* in response to climate change in the southern Africa region. As a result, this study aims to use Maxent to map the current distribution of *C. mopane*, and predict new distribution areas in the coming decades due to climate change under different representative concentration pathways (RCPs).

## 2. Materials and methods

### 2.1 Occurrence records

The distribution of *C. mopane* used in the present study was obtained from two sources: The Global Biodiversity Information Facility (GBIF, http://www.gbif.org/) (759 records) and Zambia flora (122 records). GBIF platform provides basic data on biodiversity; we searched the keyword “Colomophosum mopane” on GBIF, accessed on 12^th^ May, 2020 and downloaded the data. We conducted initial filtering of the data by removing duplicated records, followed by spatial rarefying of the data performed on R package “spThin” v. 0.1.0 (Aiello□Lammens et al., 2015) to reduce the autocorrelation between the points at each grid cell of 102 km. The remaining 367 records were used in the subsequent analyses (Figure 1).

### 2.2 Climate data and clipping

We obtained the climate data from the Worldclim database version 1.4 (Hijmans, 2005, http://www.worldclim.org). Nineteen bioclimatic variables and an elevation layer at a resolution of 2.5 arc-mic were used. To reduce collinearity, the 19 bioclimatic variables were subjected to Pearson’s correlation at a threshold of 0.8 (Graham, 2003), implemented in ENMTools package in R, using the function *raster.cor.matrix* (Warren et al., 2019). ArcGIS was used to clip the study area, including the known ranges and the potential distribution regions in Southern Africa. Eventually, eight variables were selected as representative of climate factors.

### 2.3 Model building and evaluation of SDM performance

The maximum entropy algorithm implemented in Maxent v3.3k was used to develop the current SDMs for *C. mopane* by allowing the transformations of covariates utilizing the software’s default parameters, except the following: number of background points = 10^^4^, and the number of iterations = 5000. Eighty percent of the localities for model training, and the remaining 20% for testing. The model validation involved conducting 10 replicated run models for *C*. *mopane*, applying the threshold rule of maximum training sensitivity plus specificity (MTSS). MTSS is recommended as a conservative approach that minimizes commission and omission errors (Guisan et al., 2017; Liu et al., 2016). Jackknife tests were used on each of the 10 replicated models to assess the most important variables contributing to Maxent’s final model (Phillips et al., 2006). The performance of models was evaluated by the area under the curve (AUC) values of the receiving operator curve (ROC) (Mas et al., 2013). The AUC statistics of the receiver operating characteristics (ROC) were assessed to evaluate the model performance (Phillips and Dudik, 2008). AUC values were examined using the test points. AUC values less than 0.8 indicate poor performance of the model, AUC values between 0.8-0.9 moderate performance, while AUC values above 0.9 are considered excellent (Thuiller et al., 2005). MTSS threshold was used to convert the models with probability values to binary (presence/absence). The presence of *C. mopane* was projected geographically by driving the probability of occurrence in four categories as follows: values below MTSS threshold no data, “low” threshold–0.3, “medium” 0.3–0.6, and “high” □0.6. The logistic output was used to generate the final models, where the MTSS was used to define the presence-absence binary data.

To assess the probable future distribution in *C. mopane* ranges, we utilized the climate projections from the Coupled Model Intercomparison Project Phase 5 (CMIP5) (Collins et al., 2014). We considered three RCP scenarios for the CCSM4 model (Gent et al., 2011). The stringent mitigation scenario RCP 2.6 aims to keep global warming under 2.0°C under pre-industrial temperatures by 2010 (Collins et al., 2014). Under the intermediate scenarios, the global mean surface temperatures are projected to rise by 1.5-3.2 °C and the CO2 concentrations to 850 ppm in RCP 4.5. For the RCP8.5 pessimistic scenario, global mean surface temperatures will possibly increase in by 2.6-4.8 °C, while the concentration of CO2 will approximately be 1,350 ppm by 2100.

## 3. Results

### 3.1 Variable Importance and Climatic Preference

Eight variables were retained after the correlation analyses following their multicollinearity. The model tunings test for *C. mopane* produced satisfactory outcomes for the current and future scenarios with high AUC scores = 0.941-0.942 (Table 1). These results signified the high reliability of the models in habitat suitability modeling for *C. mopane* in southern Africa. The most significant variables for the current distribution of *C. mopane* were Solar radiation (relative contribution: 41.8%), Bio12 (annual precipitation - relative contribution: 16.9%)), and Bio7 (annual temperature range - relative contribution: 15.9%), contributing to 74.6% of the maxent model (Table 2). In addition, for all the future scenarios, the same variables remained the most influential variables limiting mopane distribution. Jackknife tests showed that when used in isolation, Solar radiation, Bio12 (annual precipitation), and Bio7 (annual temperature range) had the highest gain and were considered the most informative (Fig 2). When solar radiation was omitted, the gain was reduced the most, indicating that this variable had most information absent in other variables.

**Table 1.** MaxEnt Model performance for *C. mopane*

| Period | AUCtraining | AUCtest | MTSS |
| --- | --- | --- | --- |
| Current | 0.950 | 0.941 | 0.1927 |
| RCP 2.6 2050 | 0.949 | 0.942 | 0.1830 |
| RCP 4.5 2050 | 0.950 | 0.941 | 0.1879 |
| RCP 8.5 2050 | 0.949 | 0.942 | 0.1984 |
| RCP 2.6 2070 | 0.949 | 0.942 | 0.1860 |
| RCP 4.5 2070 | 0.949 | 0.942 | 0.1808 |
| RCP 8.5 2070 | 0.948 | 0.942 | 0.2075 |

AUC, Area Under Curve; MTSS, Maximum training sensitivity plus specificity

**Table 2.** Contributions (%) of the variables to the distribution of *C. mopane* according to Maxent Modeling (bold values are the most important variables)

| Variable | Description | Current | 2050 |  |  | 2070 |  |  |
| --- | --- | --- | --- | --- | --- | --- | --- | --- |
|  |  |  | RCP2.<br>6 | RCP4.<br>5 | RCP8.<br>5 | RCP2.<br>6 | RCP4.<br>5 | RCP8.<br>5 |
| Bio1 | Annual temperature | 5.8 | 3.5 | 3.2 | 4.5 | 3 | 5.9 | 4.3 |
| Bio5 | max temperature of warmest month | 7.7 | 6.5 | 3.8 | 5.7 | 6.8 | 8.3 | 8 |
| Bio7 | annual temperature range | **15.9** | **16.0** | **13.2** | **14.3** | **16** | **16.3** | **13.3** |
| Bio11 | mean temperature of coldest quarter | 2.7 | 2.3 | 4.3 | 2.2 | 2.6 | 2.5 | 2.1 |
| Bio12 | annual precipitation | **16.9** | **19.1** | **24.1** | **19.1** | **18.9** | **15.7** | **19.1** |
| Bio15 | precipitation seasonality | 2.5 | 1.9 | 1 | 1.4 | 1.8 | 2.0 | 1.5 |
| Bio17 | precipitation of driest quarter | 0.6 | **0.8** | 0.7 | 0.6 | 0.7 | 1.0 | 0.8 |
| Bio19 | precipitation of coldest quarter | 1.3 | 3.6 | 3.7 | 4.3 | 3.7 | 3.8 | 4.2 |
| Elevation |  | 3.3 | 4.2 | 3.6 | 5.0 | 4.4 | 3.9 | 5.6 |
| Slope |  | 1.5 | **0.6** | 0.5 | 0.7 | 0.6 | 0.9 | 0.8 |
| Solar radiation |  | **41.8** | **41.5** | **41.9** | **42.0** | **41.4** | **39.6** | **40.3** |

**Figure 2.**
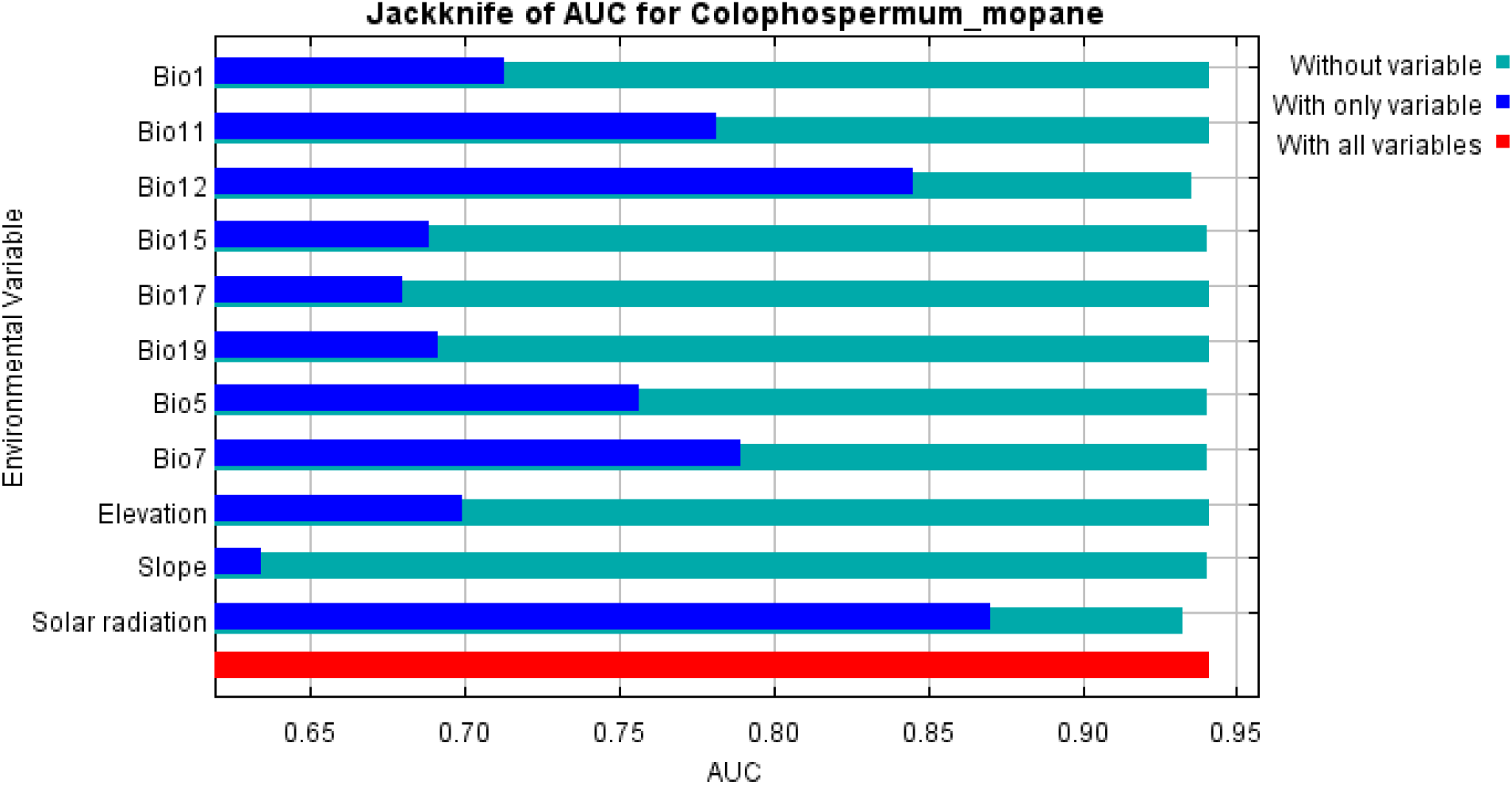
Jackknife analysis of test gain for ten runs of the current distribution model for *C. mopane*

The current distribution models showed that *C. mopane* has a potentially broader distribution in Botswana and lower elevations from approximately 15S to 25S and 11E to 35E (Fig 3). Besides, the current model’s habitat suitability revealed low suitability for *C. mopane* in Lesotho. The simulations mainly covered areas in Botswana and Zimbabwe, with central Zimbabwe, southern Mozambique, northern Namibia, and southern Zambia having the most favorable conditions. Similar high habitat suitability was also observed in northern parts of S. Africa. These high habitat suitability areas correspond with the abundant occurrences of *C. mopane*. Lower habitat suitability for *C. mopane* was observed in Madagascar, although no observations confirm its presence. Future climatic projections for *C. mopane* revealed that the pessimistic scenario RCP85 was more optimistic and highlighted the potential range expansion by 2070 compared to the intermediate pessimistic scenario RCP2.6 and RCP 4.5 (Fig 4).

**Figure 3.**
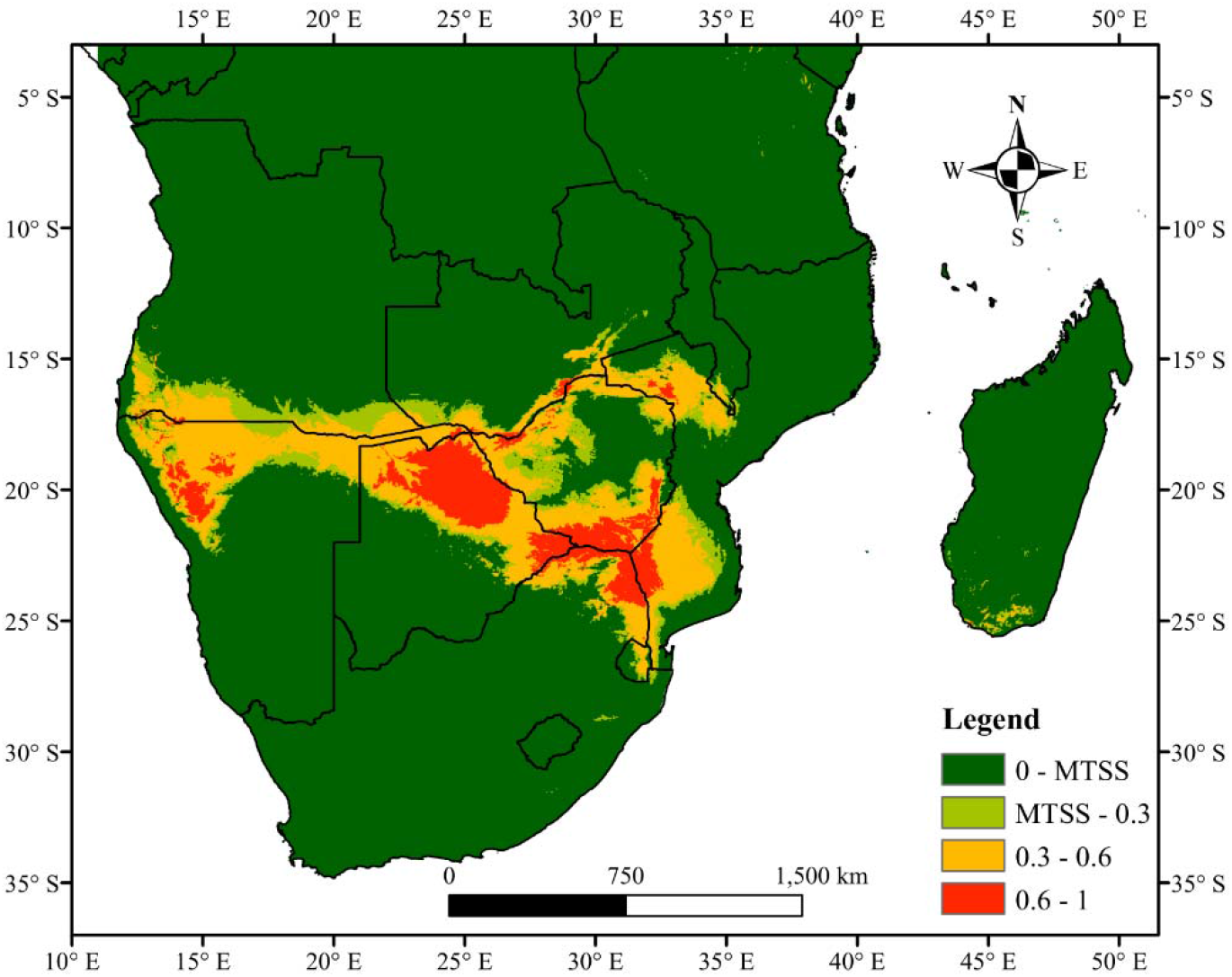
Current potential distribution map of *C. mopane*

**Figure 4.**
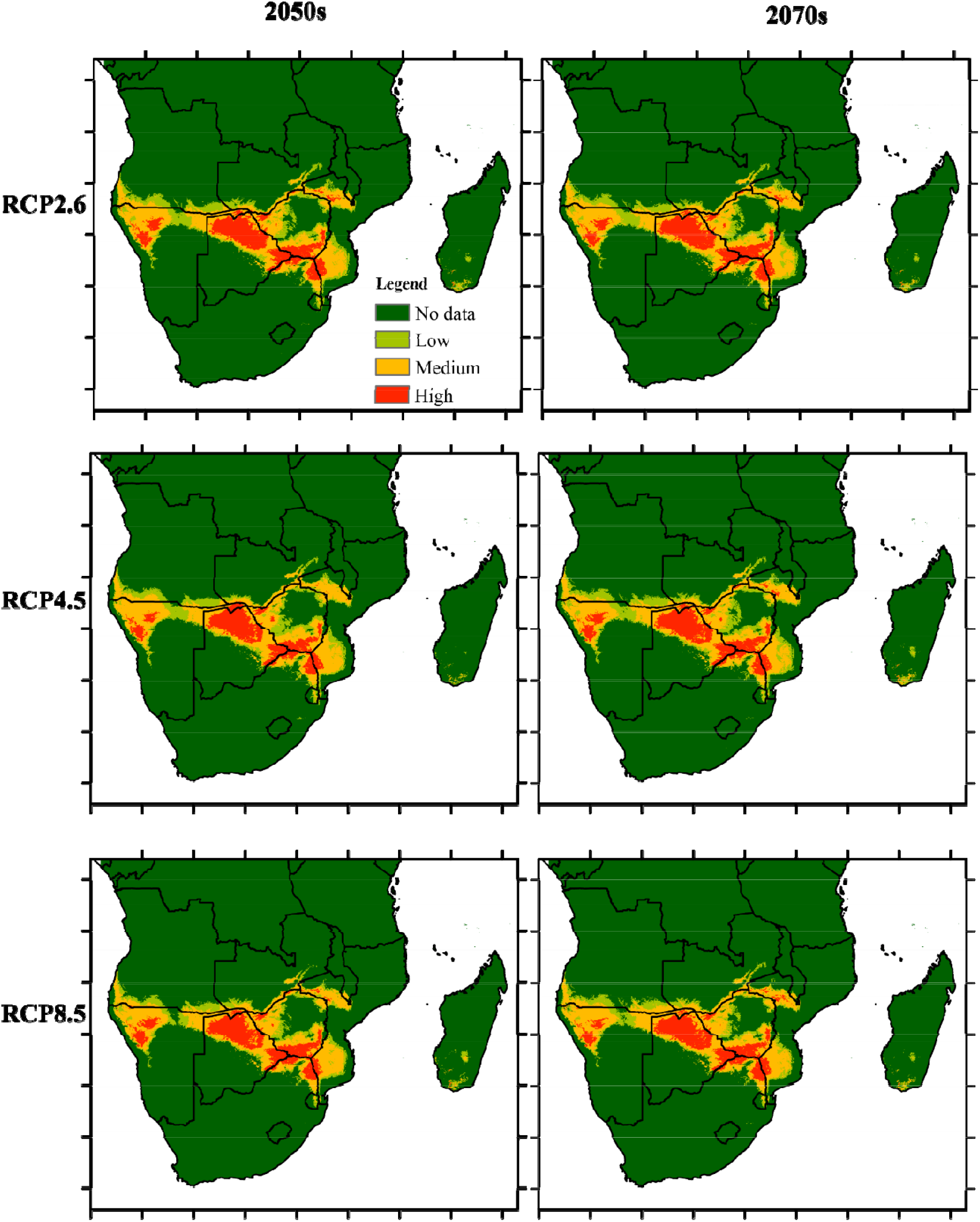
Future climatic projections of *C. mopane* in the 2050s and 2070s under different RCPs scenarios.

## Discussion

Correlative and predictive SDMs have frequently been used to produce predictions of potential species richness changes and the influence of climate change on biodiversity. Such predictions have been made for different groups of species across the planet. For example, Thuiller et al. (2005) estimated the potential loss of plant species across Europe to vary between 27% and 42% by the end of the 21st century.

As a result of temporal mismatches between species occurrence localities and current bioclimatic data, assessing SDMs in versatile geographical areas like southern Africa poses a significant challenge for model accuracy. Undeniably, previous studies assessing the distribution of mopane under different climate change scenarios have traditionally focused on a smaller portion of the mopane’s range, such as national parks, rather than its entire distribution (Stevens et al., 2014, 2018). In this study, we used the entire distribution of ranges Mopane in southern Africa, which helped us use the most amount of occurrence-environment data while avoiding errors caused by temporal mismatches.

Climate change is a major factor limiting the species’ distributions (Parmesan, 2006) and is expected to intensify in the future leading to global warming (Walther et al., 2016). In this study, SDMs were utilized to identify the current and future habitat suitability of *C. mopane* in southern Africa. Previous studies demonstrated that global climate changes had reduced the species ranges in future periods (Bellard et al., 2012; Saiz et al., 2021), with some moving polewards and to higher elevations (Lenoir et al., 2008; Parmesan & Hanley, 2015; Saiz et al., 2021). However, studies have shown that the ranges of Mopane will increase and shift westwards with increases in global warming (Stevens, 2018). Similarly, our findings have demonstrated that the habitat suitability of Mopane will shift polewards, and parts of the ranges will remain unchanged with climate change.

In estimating *C. mopane* distribution in southern Africa, our SDMs showed considerable results, supported by validation results. AUCs, such as those we obtained (> 0.941), are among the highest values for reported models and have high habitat suitability predictive capacities (Elith et al., 2010).

For the current distribution of *C. mopane*, MaxEnt projections showed that this species, in general, occurs in the warm, dry, low-lying regions. Therefore, the MaxEnt models accurately predicted the current species distribution of mopane as it tends to prefer lowland and drier habitats compared to highlands (Burgess et al., 2004; Maquia et al., 2019). Our model predicted high suitability for this tree throughout northern and eastern Botswana, southern Zimbabwe, southwestern Mozambique, and northern parts of South Africa bordering Zimbabwe and Mozambique. High habitat suitability was also observed in the northern parts of Zimbabwe bordering Zambia and northwestern Angola. These regions are characterized by extensive miombo strands and savanna ecosystems (Khavhagali, & Ligavha-Mbelengwa, 2009; Bruschi et al., 2017). The predicted habitat suitability for Mopane is consistent with previous reports that have assessed the distribution and expansion of Mopane, e.g., Kruger national park (Stevens et al., 2014, 2018). However, our model also detected the moderate habitat suitability of Mopane in southern Mozambique, southern Namibia, eastern Angola, and southern Zambia.

Solar radiation, annual temperature range, and annual precipitation contributed considerably to the models of the distribution of Mopane. Water availability and ambient temperatures are essential factors that support plant growth (Marshall, 1988). However, *C. mopane* has been observed to tolerate low nutrient conditions, moisture pressures, and even disruptions caused by fire, ability to resist drought, and browsing by large herbivores, making the species able to conquer the low-lying regions of southern Africa’s savannas (Gandiwa & Zisadza, 2011; Makhado et al., 2014).

Mopane grows in a tropical savanna climate with distinct geological and hydrological features that are ideal for the survival of mopane strands (Moura et al., 2017). As a result, environmental conditions play an essential role in the distribution of mopane. Previous research from Kruger National Park found that mopanes distribution correlated with humidity levels and temperature (Makhado et al., 2014; Stevens et al., 2014, 2018). Using the niche models, we were able to obtain concurrent research findings, demonstrating that temperature and precipitation can significantly impact mopane’s niche distribution. Notably, since precipitation is a plant growth prerequisite, it may facilitate the growth of mopane, resulting in the expansion of its natural populations.

ENM findings can also help determine a species’ physiological tolerance, which, when combined with knowledge of life history, physiological and behavioral characteristics, could help select the most plausible predictions (Escobar and Craft, 2016). Climate change is also influencing mopane populations, raising concerns about the future of woodlands in southern Africa (Stevens et al., 2021).

We choose Maxent because it consistently outperforms other predictive precision approaches, and the program is relatively user-friendly (Terribile & Diniz-Filho, 2010; Merow et al., 2013). It has been widely used to model species distributions since its publication in 2004 (Elith et al., 2011). Several experiments have been carried out to compare Maxent’s findings with other models, and Maxent has been observed to predict better areas for the use of expert landscape classification than regularized logistical regressions (Dicko et al., 2014). Maxent has also been used to forecast the spread of *C. mopane* in southern Africa (Makhado et al., 2014; Stevens et al., 2018).

The shortcomings of our research should be noted in this article; for instance, the data comes from GBIF and online databases, resulting in publication bias. Also, the precision of the geographic coordinates of mopane we obtained is limited. Lastly, the MaxEnt program does not take into account other factors influencing local adaptation and microclimates for mopane. Regrettably, we could not collect complete and up-to-date data on these factors, which should be deliberated in future research.

## 4. Conclusions

In this study, the distribution of *C. mopane* was assessed using environmental covariates and projected its future distribution under different RCPs using the current data as the baseline data. Our findings indicate that, although *C. mopane* is a semi-arid species at present living at its biological limits (Makhado et al., 2014), it may not be adversely affected by climate change, like other organisms have been shown to (Parmesan, 2006). Even so, this has not always been the scenario. In reality, certain organisms may even be able to respond to shifting local environments by phenotypic plasticity (Donelson et al., 2019). However, most plants are far more likely to shift their range and then go into extinction in response to rising temperature increases and shifts in rainfall (Williams and Blois, 2018; Dyderski et al., 2018). To introduce the most ambitious adaptation policies proposed by Yalcin and Leroux (2017), we will require precise estimates of the potential climate change impacts on biodiversity

## References

Aiello□Lammens, M. E., Boria, R. A., Radosavljevic, A., Vilela, B. & Anderson, R. P., 2015. spThin: An R package for spatial thinning of species occurrence records for use in ecological niche models. Ecography, 38(5), 541–545. https://doi.org/10.1111/ecog.01132

Austin, M.P., Van Niel, K.P., 2011. Improving species distribution models for climate change studies: variable selection and scale. J. Biogeogr. 38, 1–8. https://doi.org/10.1111/j.1365-2699.2010.02416.x

Baldwin, R. A., 2009. Use of maximum entropy modeling in wildlife research. Entropy, 11(4), 854–866.

Bellard, C., Bertelsmeier, C., Leadley, P., Thuiller, W., & Courchamp, F., 2012. Impacts of climate change on the future of biodiversity. Ecology letters, 15(4), 365–377.

Bentlage, B., Peterson, A. T., Barve, N., & Cartwright, P., 2013. Plumbing the depths: extending ecological niche modelling and species distribution modelling in three dimensions. Global Ecology and Biogeography, 22(8), 952–961.

Blanco, J.A., Ameztegui, A., Rodríguez, F., 2020. Modelling Forest Ecosystems: A crossroad between scales, techniques, and applications. Ecol. Model 425, 109030.

Bruschi, P., Urso, V., Solazzo, D., Tonini, M., & Signorini, M. A., 2017. Traditional knowledge on ethno-veterinary and fodder plants in South Angola: an ethnobotanic field survey in Mopane woodlands in Bibala, Namibe province. Journal of Agriculture and Environment for International Development (JAEID), 111(1), 105–121.

Burgess, N., Hales, J.D., Underwood, E., Dinerstein, E., Olson, D., Itoua, I., Schipper, J., Ricketts, T., Newman, K., 2004. Terrestrial ecoregions of Africa and Madagascar: a conservation assessment. Island Press, Washington.

Collins M, Knutti R, Arblaser J, Dufresne J-L, Fichefet T et al., 2014. Long-term climate change: projections, commitments and irreversibility. In: Stocker T, Qin D, Plattner G, Tignor M, Allen SK et al (eds) Climate Change 2013: The Physical Science Basis. Contribution of Working Group I to the Fifth Assessment Report of the Intergovernmental Panel on Climate Change. Cambridge University Press, Cambridge, 1029–1136

Dallas, T., Decker, R.R., Hastings, A., 2017. Species are not most abundant in the centre of their geographic range or climatic niche. Ecol. Lett. 20, 1526–1533. https://doi.org/https://doi.org/10.1111/ele.12860.

Daru, B.H., Berger, D.K., van Wyk, A.E., 2016. Opportunities for unlocking the potential of genomics for African trees. New Phytol. 210, 772–778. https://doi.org/https://doi.org/10.1111/nph.13826

Dewees, P.A., Campbell, B.M., Katerere, Y., Sitoe, A., Cunningham, A.B., Angelsen, A., Wunder, S., 2010. Managing the Miombo Woodlands of Southern Africa: Policies, Incentives and Options for the Rural Poor. J. Nat. Resour. Policy Res. 2, 57–73. https://doi.org/10.1080/19390450903350846

Dicko, A.H., Lancelot, R., Seck, M.T., Guerrini, L., Sall, B., Lo, M., Vreysen, M.J., Lefrançois, T., Fonta, W.M., Peck, S.L. and Bouyer, J., 2014. Using species distribution models to optimize vector control in the framework of the tsetse eradication campaign in Senegal. Proceedings of the National Academy of Sciences, 111(28), pp.10149–10154.

Donelson, J.M., Sunday, J.M., Figueira, W.F., Gaitán-Espitia, J.D., Hobday, A.J., Johnson, C.R., Leis, J.M., Ling, S.D., Marshall, D., Pandolfi, J.M. and Pecl, G., 2019. Understanding interactions between plasticity, adaptation and range shifts in response to marine environmental change. Philosophical Transactions of the Royal Society B, 374(1768), 20180186.

Dyderski, M. K., Paź, S., Frelich, L. E., & Jagodziński, A. M., 2018. How much does climate change threaten European forest tree species distributions? Global change biology, 24(3), 1150–1163.

Elith, J., Kearney, M., & Phillips, S., 2010. The art of modelling range□shifting species. Methods in ecology and evolution, 1(4), 330–342.

Elith, J., Phillips, S. J., Hastie, T., Dudík, M., Chee, Y. E., & Yates, C. J., 2011. A statistical explanation of MaxEnt for ecologists. Diversity and distributions, 17(1), 43–57.

Escobar, L.E., Craft, M.E., 2016. Advances and limitations of disease biogeography using ecological niche modeling. Front. Microbiol. 7, 1174.

Fourcade, Y., Engler, J. O., Rödder, D., & Secondi, J., 2014. Mapping species distributions with MAXENT using a geographically biased sample of presence data: a performance assessment of methods for correcting sampling bias. PloS one, 9(5), e97122.

Gandiwa, E., & Zisadza, P., 2011. Wildlife management in Gonarezhou National Park, Southeast Zimbabwe: climate change and implications for management. Nature and Faune, 25(1), 101–110.

Graham, M. H., 2003. Confronting multicollinearity in ecological multiple regression. Ecology, 84(11), 2809–2815.

Gent, P.R., Danabasoglu, G., Donner, L.J., Holland, M.M., Hunke, E.C., Jayne, S.R., Lawrence, D.M., Neale, R.B., Rasch, P.J., Vertenstein, M. & Worley, P.H., 2011. The community climate system model version 4. Journal of climate, 24(19): 4973–4991. https://doi.org/10.1175/2011JCLI4083.1

Guisan, A., & Thuiller, W., 2005. Predicting species distribution: offering more than simple habitat models. Ecology letters, 8(9), 993–1009.

Guisan, A.; Thuiller, W.; Zimmermann, N.E., 2017. Habitat suitability and distribution models: with applications in R; Cambridge University Press; ISBN 0521765137.

Handa, A.K., Chavan, S.B., Sirohi, C., Rizvi, R.H., 2020. Importance of agroforestry systems in carbon sequestration, in: Proceedings of the National Agroforestry Symposium.

Hartnett, D. C., Ott, J. P., Sebes, K., & Ditlhogo, M. K., 2012. Coping with herbivory at the juvenile stage: responses to defoliation and stem browsing in the African savanna tree Colophospermum mopane. Journal of Tropical Ecology, 161–169.

Hijmans, R. J., Cameron, S. E., Parra, J. L., Jones, P. G., & Jarvis, A., 2005. Very high resolution interpolated climate surfaces for global land areas. International Journal of Climatology: A Journal of the Royal Meteorological Society, 25(15), 1965–1978.

Kennedy, A.D., Potgieter, A.L.F., 2003. Fire season affects size and architecture of Colophospermum mopane in southern African savannas. Plant Ecol. 167, 179–192. https://doi.org/10.1023/A:1023964815201

Khavhagali, V. P., & Ligavha-Mbelengwa, M. H., 2009. Colophospermum mopane woodlands of Makuya Nature Reserve, Limpopo Province.

Krug, J.H.A., 2017. Adaptation of Colophospermum mopane to extra-seasonal drought conditions: site-vegetation relations in dry-deciduous forests of Zambezi region (Namibia). For. Ecosyst. 4, 25. https://doi.org/10.1186/s40663-017-0112-0.

Langley, J., Van der Westhuizen, S., Morland, G., van Asch, B., 2020. Mitochondrial genomes and polymorphic regions of Gonimbrasia belina and Gynanisa maja (Lepidoptera: Saturniidae), two important edible caterpillars of Southern Africa. Int. J. Biol. Macromol. 144, 632–642. https://doi.org/https://doi.org/10.1016/j.ijbiomac.2019.12.055

Liu, C.; Newell, G.; White, M., 2016. On the selection of thresholds for predicting species occurrence with presence□only data. Ecol. Evol., 6, 337–348.

Lenoir, J., Gégout, J.-C., Marquet, P.A., De Ruffray, P., Brisse, H., 2008. A significant upward shift in plant species optimum elevation during the 20th century. Science (80-.). 320, 1768–1771.

Lewis, J.S., Farnsworth, M.L., Burdett, C.L., Theobald, D.M., Gray, M., Miller, R.S., 2017. Biotic and abiotic factors predicting the global distribution and population density of an invasive large mammal. Sci. Rep. 7, 44152. https://doi.org/10.1038/srep44152

Longmore, R., Busby, J.R., 1986. Atlas of elapid snakes of Australia. Australian Govt. Pub. Service.

Makhado, R.A., Mapaure, I., Potgieter, M.J., Luus-Powell, W.J., & Saidi, A.T., 2014. Factors influencing the adaptation and distribution of *Colophospermum mopane* in southern Africa’s mopane savannas - A review. Bothalia-African Biodiversity & Conservation, 44(1), 1–9.

Maquia, I., Catarino, S., Pena, R.A., Brito, R.A.D., Ribeiro, S.N., Romeiras, M.M., Ribeiro-Barros, I.A., 2019. Diversification of African Tree Legumes in Miombo–Mopane Woodlands. Plants. https://doi.org/10.3390/plants8060182

Marshall, R. H., 1988, February. Environmental factors affecting plant productivity in. In Fort Keogh research symposium (Vol. 1, pp. 27–32).

Mas J-F, Soares Filho B, Pontius RG, Farfán Gutiérrez M, Rodrigues H., 2013. A suite of tools for ROC analysis of spatial models. ISPRS Int J Geo-Inf 2:869–887. https://doi.org/10.3390/ijgi2030869

Merow, C., Smith, M. J., & Silander Jr, J. A., 2013. A practical guide to MaxEnt for modeling species’ distributions: what it does, and why inputs and settings matter. Ecography, 36(10), 1058–1069.

Mlambo, D., Nyathi, P., Mapaure, I., 2005. Influence of Colophospermum mopane on surface soil properties and understorey vegetation in a southern African savanna. For. Ecol. Manage. 212, 394–404. https://doi.org/https://doi.org/10.1016/j.foreco.2005.03.022

Mojeremane, W., & Lumbile, A. U., 2005. The characteristics and economic values of Colophospermum mopane (Kirk ex Benth.) J Léonard in Botswana. Pakistan Journal of Biological Sciences, 8(5), 781–784.

Moura, I., Maquia, I., Rija, A. A., Ribeiro, N., & Ribeiro-Barros, A. I., 2017. Biodiversity studies in key species from the African mopane and miombo woodlands. Genetic Diversity; Bitz, L., Ed.; IntechOpen: London, UK, 91–109. https://doi.org/10.5772/66845

Pachauri, R.K., Allen, M.R., Barros, V.R., Broome, J., Cramer, W., Christ, R., Church, J.A., Clarke, L., Dahe, Q., Dasgupta, P., 2014. Climate change 2014: synthesis report. Contribution of Working Groups I, II and III to the fifth assessment report of the Intergovernmental Panel on Climate Change. Ipcc.

Parmesan, C., 2006. Ecological and evolutionary responses to recent climate change. Annu. Rev. Ecol. Evol. Syst., 37, 637–669.

Parmesan, C., Hanley, M.E., 2015. Plants and climate change: complexities and surprises. Ann. Bot. 116, 849–864.

Phillips, S. J., Dudík, M., & Schapire, R. E., 2006. A maximum entropy approach to species distribution modeling. In Proceedings of the twenty-first international conference on Machine learning (p. 83).

Phillips, S. J., 2008. Transferability, sample selection bias and background data in presence-only modelling: a response to Peterson et al.(2007). Ecography, 31(2), 272–278.

Phillips, S. J., & Dudík, M., 2008. Modeling of species distributions with Maxent: new extensions and a comprehensive evaluation. Ecography, 31(2), 161–175.

Rosenstock, T.S., Dawson, I.K., Aynekulu, E., Chomba, S., Degrande, A., Fornace, K., Jamnadass, R., Kimaro, A., Kindt, R., Lamanna, C., 2019. A planetary health perspective on agroforestry in Sub-Saharan Africa. One Earth 1, 330–344.

Saiz, H., Dainese, M., Chiarucci, A., & Nascimbene, J., 2021. Networks of epiphytic lichens and host trees along elevation gradients: Climate change implications in mountain ranges. Journal of ecology, 109(3), 1122–1132.

Stevens, N., Archibald, S.A., Bond, W.J., 2018. Transplant Experiments Point to Fire Regime as Limiting Savanna Tree Distribution. Front. Ecol. Evol..

Stevens, N., Swemmer, A.M., Ezzy, L., Erasmus, B.F.N., 2014. Investigating potential determinants of the distribution limits of a savanna woody plant: *Colophospermum mopane*. J. Veg. Sci. 25, 363–373. https://doi.org/10.1111/jvs.12098

Stevens, N., 2021. What shapes the range edge of a dominant African savanna tree, Colophospermum mopane? A demographic approach. Ecology and Evolution.

Terribile, L. C., & Diniz-Filho, J. A. F., 2010. How many studies are necessary to compare niche-based models for geographic distributions? Inductive reasoning may fail at the end. Brazilian Journal of Biology, 70(2), 263–269.

Thuiller, W., Richardson, D. M., Pyšek, P., Midgley, G. F., Hughes, G. O., & Rouget, M., 2005. NicheLbased modelling as a tool for predicting the risk of alien plant invasions at a global scale. Global change biology, 11(12), 2234–2250.

Walther, G.R., Post, E., Convey, P., Menzel, A., Parmesan, C., Beebee, T.J., Fromentin, J.M., Hoegh-Guldberg, O. and Bairlein, F., 2002. Ecological responses to recent climate change. Nature, 416(6879), pp.389–395.

Warren DL, Matzke N, Cardillo M, Baumgartner J, Beaumont L et al., 2019. ENMTools R package (software package). Species Space. https://doi.org/10.5281/zenodo.3268814

Williams, J. E., & Blois, J. L., 2018. Range shifts in response to past and future climate change: Can climate velocities and species’ dispersal capabilities explain variation in mammalian range shifts? Journal of Biogeography, 45(9), 2175–2189.

Yalcin, S., & Leroux, S. J., 2017. Diversity and suitability of existing methods and metrics for quantifying species range shifts. Global Ecology and Biogeography, 26(6), 609–624

